# Correlated Segments of Intrinsically Disordered Proteins as Drivers of Homotypic Phase Separation

**DOI:** 10.1101/2025.04.06.647444

**Authors:** Huan-Xiang Zhou

## Abstract

Many studies have suggested that amino acid composition, not their positions along the sequence, is the determinant of phase separation of intrinsically disordered proteins (IDPs). In particular, aromatic amino acids and Arg have been identified as major drivers. Here I underscore the importance of the positions of amino acids along the sequence in phase separation. Specifically, clusters of interaction-prone amino acids, including Trp and Arg, along the sequence form correlated segments, and these correlated segments, rather than individual residues, drive the phase separation of many IDPs. Correlated segments manifest themselves as stretches of residues that span major peaks in the backbone ^15^N NMR transverse relaxation rates and can be predicted by a sequence-based method called SeqDYN (https://zhougroup-uic.github.io/SeqDYNidp/). Inter-chain interactions between individual residues may be too transient, but those between correlated segments involve multiple residues can provide the strengths required for phase separation. Indeed, sequence motifs revealed by NMR and other techniques as important for phase separation frequently map to SeqDYN-predicted correlated segments. These include residues G624–R626, G638–R640, and R660–Q666 of CAPRIN1, residues R21-G30 of LAF-1, and residues Q9-P21 of FUS. SeqDYN presents a sequence-based method for identifying motifs that drive phase separation of IDPs.

## Introduction

Surface attractive patches play critical roles in the liquid-liquid phase separation (LLPS) of folded proteins and proteins containing folded domains ^1-3^. Surface sites on folded domains such as SH3 and SUMO are also important in driving heterotypic phase separation, engaging with cognate interaction motifs (proline-rich or SUMO-interacting) in intrinsically disordered proteins (IDPs) ^3-5^. For IDPs, the pre-organization of residues distant along the sequence into clusters akin to surface patches or surface sites on folded domains are likely to be very rare. Instead, stretches of interaction-prone residues may form correlated segments. The general question in the present study is whether correlated segments of IDPs can be the drivers of homotypic phase separation.

Many IDPs readily undergo homotypic phase separation due to distinct advantages conferred by their disordered nature, including the potential for all residues, not just surface residues as in folded proteins, to form intermolecular interactions ^6,7^. How the amino-acid sequence of an IDP codes for its propensity for and other properties of phase separation is a fundamental, but still open question. Mutational effects have amply demonstrated that the importance of aromatic amino acids and Arg in driving the phase separation of IDPs ^8-15^. A view has emerged that the composition of amino acids, not their positions along the sequence, is the major determinant for IDP phase separation ^9,10,14,16^. In particular, empirical formulas were proposed to explain the threshold (or saturation) concentrations of phase separation in terms of compositional parameters such as the numbers of aromatic and Arg residues ^10,14^. Properties solely derived from amino-acid composition have been used to predict or rationalize phase-separation propensities ^17-19^.

The composition-based view implies that LLPS-driving amino acids (“stickers”) act independently of each other. A prime example conforming to this scenario is the low-complexity domain of hnRNPA1 (A1-LCD) ^12^. This IDP has a high aromatic content (14% of the sequence); these aromatic residues are more uniformly distributed along the sequence than in 99.99% of randomly generated sequences. Repositioning the aromatic residues into clusters of three or four along the sequence resulted in amorphous aggregates instead of liquid droplets. It appears that, due to the high aromatic content in A1-LCD, pairwise attraction between aromatic residues is sufficient for LLPS. Clustering of these aromatic residues can increase the drive for phase separation, but at the risk of producing overly strong attraction as to resulting in solid-like aggregates instead of liquid droplets.

The importance of the positions of amino acids along the sequence in IDP phase separation is demonstrated by the effects of charge shuffling. Segregation of opposite charges into charge blocks strengthens electrostatic attraction and promotes phase separation, as demonstrated by coarse-grained simulations ^20^ and by experimental effects of charge shuffling to either decrease ^8^ or increase charge blockiness ^13,21^. In comparison, sequence shuffling in a computational study of the FUS LCD suggested that stretches of 2-3 hydrophobic residues are crucial for driving the phase separation of this IDP ^22^. A parameter describing long-range hydrophobic patterning did not show correlation with phase-separation propensity in shuffling exercises ^23^. A middle ground, with residue-residue interaction parameters along two sequences (of the same IDP for homotypic LLPS) averaged over a 13-residue sliding window, has shown promise in predicting sequence motifs driving the phase separation of DDX4 ^24^.

Sequence sub-regions, motifs, and residues that are essential for phase separation have been identified for many IDPs. Using deletion constructs, Tari et al. ^25^ and Yu et al. ^26^ found that an Arg/Ser-rich (RS) region and a coiled-coiled sub-region, respectively, drive the LLPS of the splicing factor U2AF^65^ and the C-terminal intrinsically disordered region (IDR) of the yes-associated protein (YAP). Coarse-grained simulations of Schuster et al. ^13^ revealed high inter-chain contact probabilities for residues 21-30 in the LAF-1 intrinsically disordered RGG domain; their importance in LLPS was confirmed by a significant decrease in cloud-point temperature upon deletion. Various NMR experiments have been performed to uncover interacting residues. Intermolecular nuclear Overhauser effects (NOEs) were measured in the dense phase after phase separation; in most cases it was only possible to assign the cross peaks to amino-acid types ^27-31^ but for the CAPRIN1 C-terminal IDR, residue-specific assignments were made, revealing residues 624–626, 638–640, and 660–666 as critical for phase separation ^32^. Intermolecular paramagnetic relaxation enhancements (PREs) revealed high inter-chain interaction propensities for residues around Ala16 in FUS LCD ^28^. Lastly, backbone ^15^N transverse relaxation rates measured below the threshold concentration were used to dissect drivers of IDP phase separation ^12,15,32-34^.

Backbone ^15^N transverse relaxation rates, denoted as *R*_2_, are elevated when residues are part of a “correlated” segment rigidified by either local inter-residue interactions or residual secondary structure ^35,36^. As suggested previously ^15,32,37,38^, in the dense phase after phase separation, each IDP chain is surrounded by other chains, and intra-chain interactions give way to inter-chain interactions. Therefore correlated segments are very likely to participate in inter-chain interactions and thereby drive phase separation. We have previously developed a sequence-based method, SeqDYN, to predict *R*_2_ profiles of IDPs ^37^. Here I show that LLPS-driving sequence motifs frequently map to segments with elevated values in SeqDYN-predicted *R*_2_ profile. These results justify both the notion that clusters of interaction-prone amino acids along the sequence, rather than the amino acids separately, drive the phase separation of many IDPs and the use of SeqDYN for identifying such sequence motifs.

## Results

### The SeqDYN *q* parameters correlate with phase-separation propensities of tetrapeptides

SeqDYN ^37^ was designed to predict the backbone ^15^N transverse relaxation rates, *R*_2_, which are sensitive to motions in the 1-10 ns range and thereby affected by local interactions and residual secondary structures ^36^. SeqDYN assumes that, for the *R*_2_ value of residue *n*, each residue *i* contributes a factor *f*(*i*; *n*), leading to

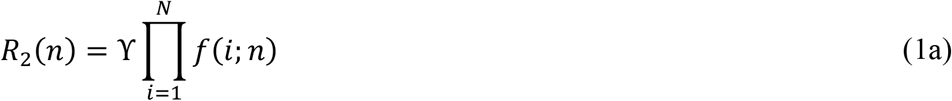

where *N* is the total number of residues in the IDP and ϒ is a uniform scale parameter. The contributing factor *f*(*i*; *n*) has an amplitude *q*(*i*) that is determined by the amino-acid type of residue *i* and attenuates with increasing sequence distance, *s* = |*i* − *n*|, from residue *n*:

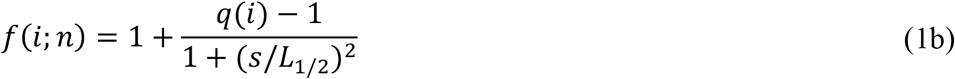

where *L*_1/2_ is the correlation half-length, i.e., the sequence distance at which *f*(*i*; *n*) is halfway between the amplitude *q*(*i*) and the baseline value of 1. The values of the 20 *q* parameters (one for each amino-acid type) and *L*_1/2_ were obtained by optimizing SeqDYN predictions against the measured *R*_2_ values for 45 IDPs.

The *q* parameters, displayed in Figure 1a, range from 1.39 for Trp to 0.98 for Gly. Notably, Trp, Tyr, and Arg, which are often cited as drivers of phase separation, are among the four amino acids with the highest *q* values. The two remaining aromatic amino acids, Phe and His, and two long-branched aliphatic amino acids, Ile and Leu, complete the top-seven subset. *q* captures the intrinsic tendency of an amino acid to interact with other amino acids, in the context of a single chain for *R*_2_ prediction. The makeup of the top-seven subset suggests the importance of π-π, cation-π, and hydrophobic interactions in elevating the *R*_2_ rates of IDPs at the single-chain level. As intra-chain interactions convert to inter-chain interactions during homotypic phase separation, *q* may capture the propensity of an amino acid to form such inter-chain interactions and hence the drive for phase separation. This contention is supported by a strong correlation between amino acid-amino acid contacts in a single chain and those in the dense phase calculated in atomistic simulations ^15^. Here I directly test *q* against the phase-separation propensities of tetrapeptides of the form XXssXX (Figure 1b), where X denotes any amino acid and ss represents a disulfide bond along the backbone ^39^. Zhang et al. ^40^ measured the threshold (or saturation) concentrations (*C*_th_) of eight tetrapeptides with X = W, F, L, I, M, V, A, and P. A low *C*_th_ represents a high phase-separation propensity. At their threshold concentrations, the peptides were observed in various material states, including droplets (X = F and L; Figure 1c), aggregates (X = W), gels (X = I), and amorphous dense liquids (X = M, V, A, and P). There is a strong correlation between − In(*C*_th_) and *q*, with a Pearson correlation coefficient (*r*) of 0.75 (Figure 1d).

**Figure 1.**
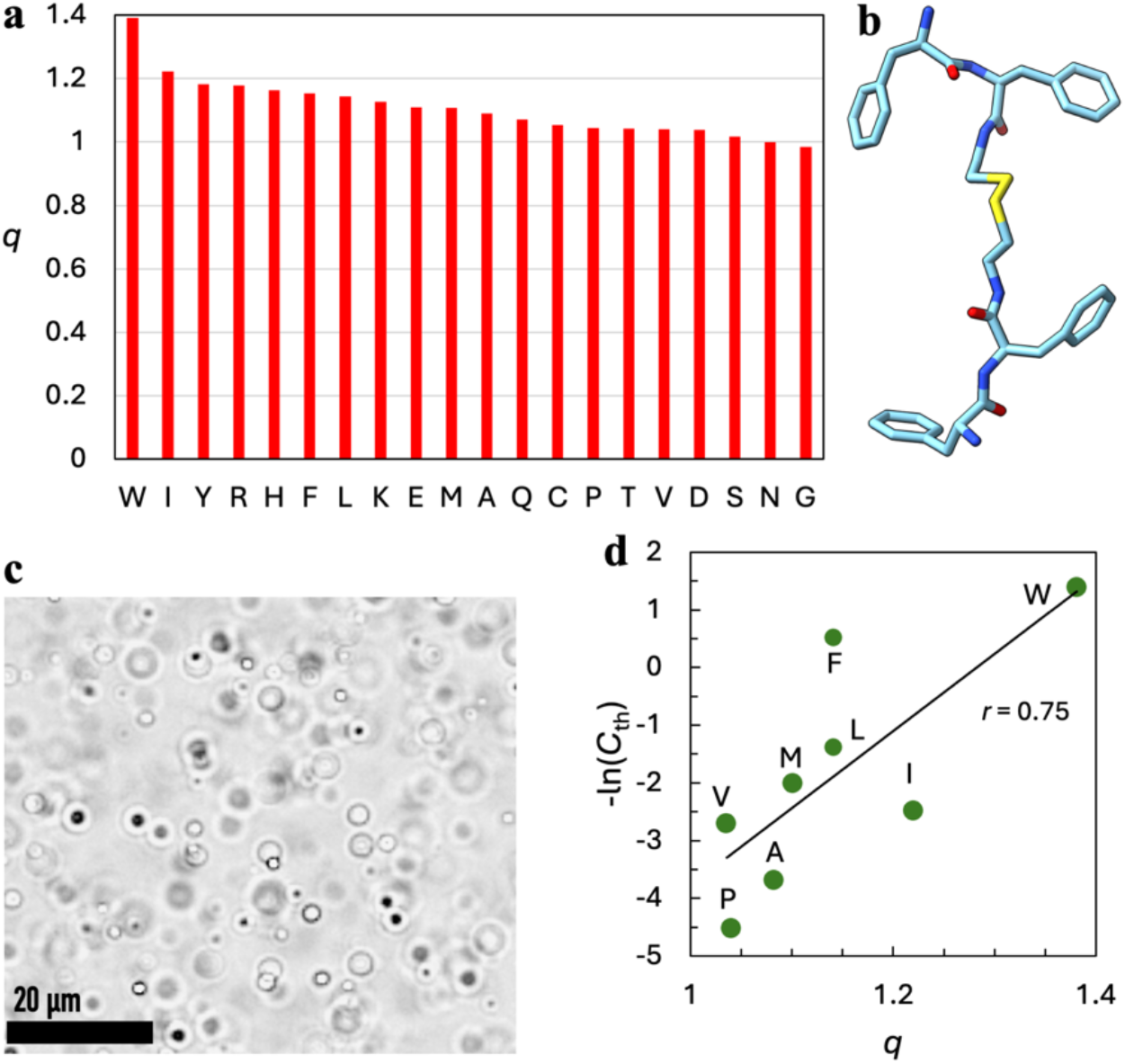
The *q* parameters predict well the phase-separation propensities of XXssXX tetrapeptides. (a) Values of *q* parameters, from Qin and Zhou ^37^. (b) Structure of FFssFF. (c) Brightfield image of droplets formed by FFssFF, from Zhang et al. ^40^. (d) Correlation between − In(*C*_th_) and *q*.

The correlation half-length *L*_1/2_ has a value of 5.6, meaning that residues 5 to 6 positions downstream and upstream can still influence the *R*_2_ value of a given residue. The influence arises from residue-residue interactions. A stretch of residues with high *q* values can form a correlated segment, where the residues engage with each other via a variety of interactions including π-π, cation-π, and salt bridges, as observed in atomistic simulations ^35,36^. Correlated segments can be identified from major peaks in the *R*_2_ profile along the sequence. The main point of this paper is that correlated segments on different IDP chains interact with each other and drive phase separation.

### CAPRIN1: three LLPS-driving motifs exhibit major peaks in the *R*_2_ profile

Based on intermolecular NOEs, Kim et al.^32^ identified three motifs, G_624_YR_626_, G_638_YR_640_, and R_660_DYSGYQ_666_, in CAPRIN1 C-terminal IDR (residues 607-709) as drivers of its phase separation. These authors also recognized that these motifs show elevated *R*_2_ rates. In the SeqDYN-predicted *R*_2_ profile, these three motifs are at the highest peaks (regions shaded in yellow in Figure 2a and residues with sidechains displayed in Figure 2b). This outcome suggests that SeqDYN can used to predict LLPS-driving motifs in IDPs. The elevated *R*_2_ rates of the motifs are firstly due to the high-*q* amino acids like Tyr and Arg within these motifs but are also contributed by high-*q* amino acids downstream and upstream of these motifs, such as Y636 and F643 around the second motif and F656, R667, and Y670 around the third motif.

**Figure 2.**
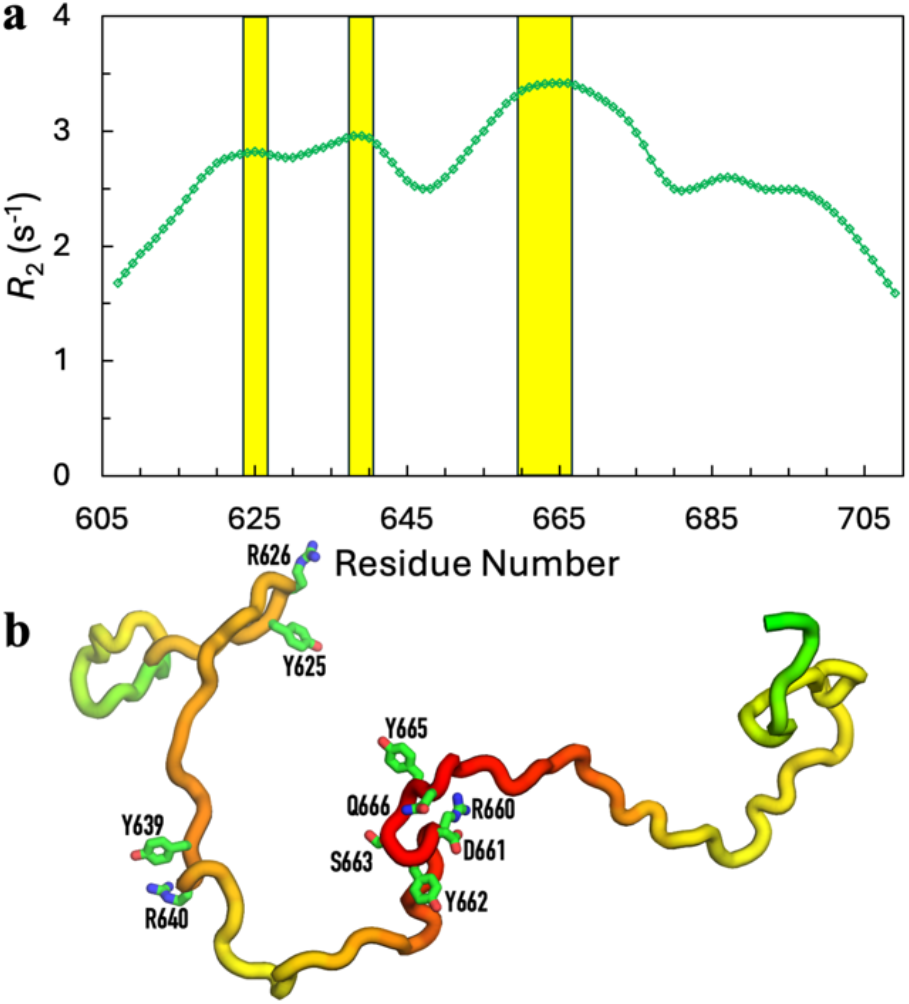
SeqDYN predicts the sequence motifs driving the phase separation of CAPRIN1. (a) *R*_2_ profile. Yellow shading highlights three major peaks, spanned by G624–R626, G638–R640, and R660–Q666 that were identified by NOESY ^32^. (b) *R*_2_ displayed as a color spectrum (green-yellow-orange-red) rendered on a representative CAPRIN1 structure. Sidechains are displayed for motifs highlighted in (a).

Kim et al. mutated three consecutive residues in these motifs into ASA, and found reduced LLPS propensities. Correspondingly, the *R*_2_ rates in and around the mutation sites, A_624_SA_626_, A_638_SA_640_, and A_660_SA_662_, showed significant decreases. SeqDYN predicts well these changes in *R*_2_ profiles (Figure S1), and thus captures the reduced drive for LLPS in the mutants. Interestingly, the measured *R*_2_ difference profiles, *R*_2_(WT) − *R*_2_(mutant), have a Lorentzian shape with a slowly decaying tail, precisely as modeled by SeqDYN [eq (1b)]. The accurate prediction of the *R*_2_ difference profiles between wild-type (WT) CAPRIN1 and the three mutants directly validates the value selected for the correlation half-length *L*_1/2_.

### LAF-1: an interaction-prone segment promotes LLPS

Coarse-grained simulations of Schuster et al. ^13^ showed high inter-chain contact probabilities for residues R_21_YVPPHLRGG_30_ in the LAF-1 RGG domain (residues 1-168). Residues R_21_YVPPHLR_28_ are also highly conserved in the DDX3 family of RNA helicases and constitute the binding motif for the eukaryotic translation initiation factor 4E. Deletion of residues 21-30 resulted in a 12 °C decrease in cloud-point temperature.

Figure 3 displays the SeqDYN-predicted *R*_2_ profile, which exhibits the highest peak within residues 21-30 (yellow shading in panel *a* and sidechains in panel *b*). So here again a correlated segment plays a major role in driving phase separation. In this segment, a half of the residues are in the top-7 subset for *q* values. As controls, Schuster et al. ^13^ deleted 10-residue segments at two other positions, 82-91 and 101-110, and a “much more modest reduction” in cloud-point temperature. In line with this experimental observation, these two segments each contain a local minimum in *R*_2_.

**Figure 3.**
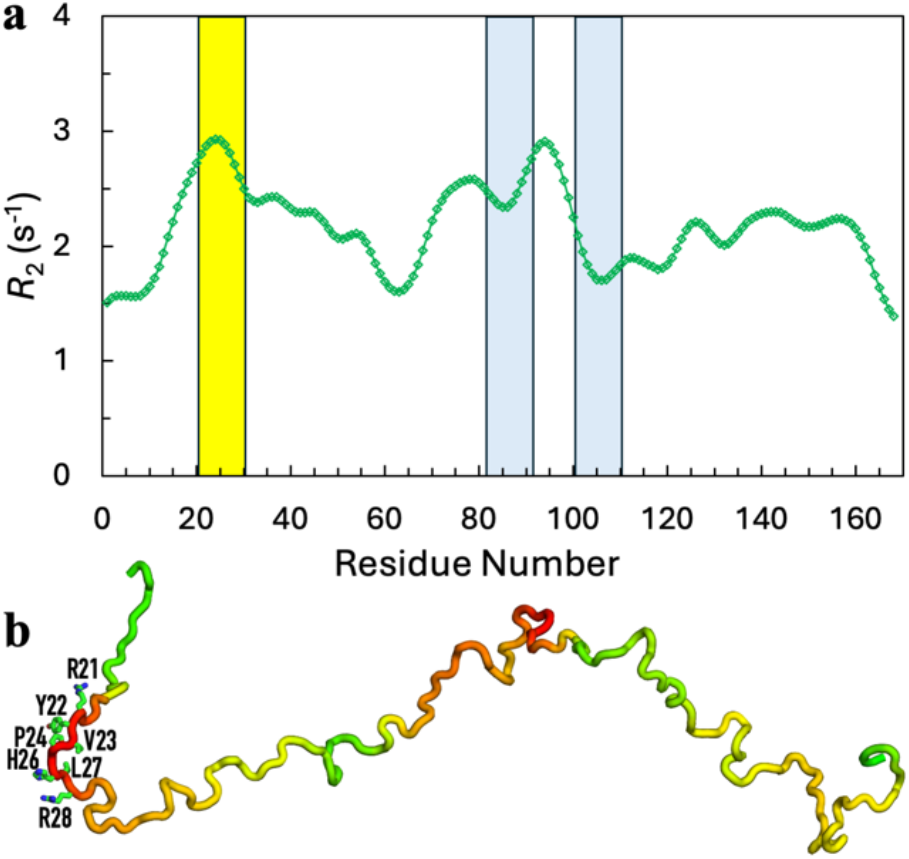
SeqDYN predicts the sequence motif driving the phase separation of LAF-1. (a) *R*_2_ profile. Yellow shading highlights the highest peak, spanned by R21-G30 that was identified by coarse-grained simulations and validated by deletion effects ^13^. Shaded in light blue are two regions introduced as negative controls in the latter study; they span local minima of the *R*_2_ profile. (b) *R*_2_ displayed as a color spectrum (green-yellow-orange-red) rendered on a representative LAF-1 structure. Sidechains are displayed for the motif highlighted in yellow in (a).

### FUS: an N-terminal segment preferentially forms inter-chain interactions in the dense phase

Murthy et al. ^28^ reported intermolecular NOEs in the dense phase of FUS LCD (residues 1-163), showing that all the major amino-acid types in the sequence participate in intermolecular interactions. In addition, PREs (denoted as *Γ*_2_) were observed throughout the FUS LCD. However, a spin label attached to residue 16 produced much higher PREs than those attached to residues 86 and 142, suggesting that, relative to the rest of the sequence, an N-terminal segment may be favored in forming inter-chain interactions.

Residue 16 is at the top of the highest peak in the SeqDYN-predicted *R*_2_ profile (Figure 4a). This peak is spanned by residues Q_9_ATQSYGA_16_YPTQP_21_. Although Ala16 itself is not a high-*q* residue, its *R*_2_ is elevated by two neighboring Tyr residues. Interestingly, the variations of the *R*_2_ profile along the sequence are qualitatively similar to those of the intermolecular PREs of the spin label attached to residue 16 (Figure 4b), supporting the notion that the *R*_2_ profile captures the interchain interaction propensities of individual residues.

**Figure 4.**
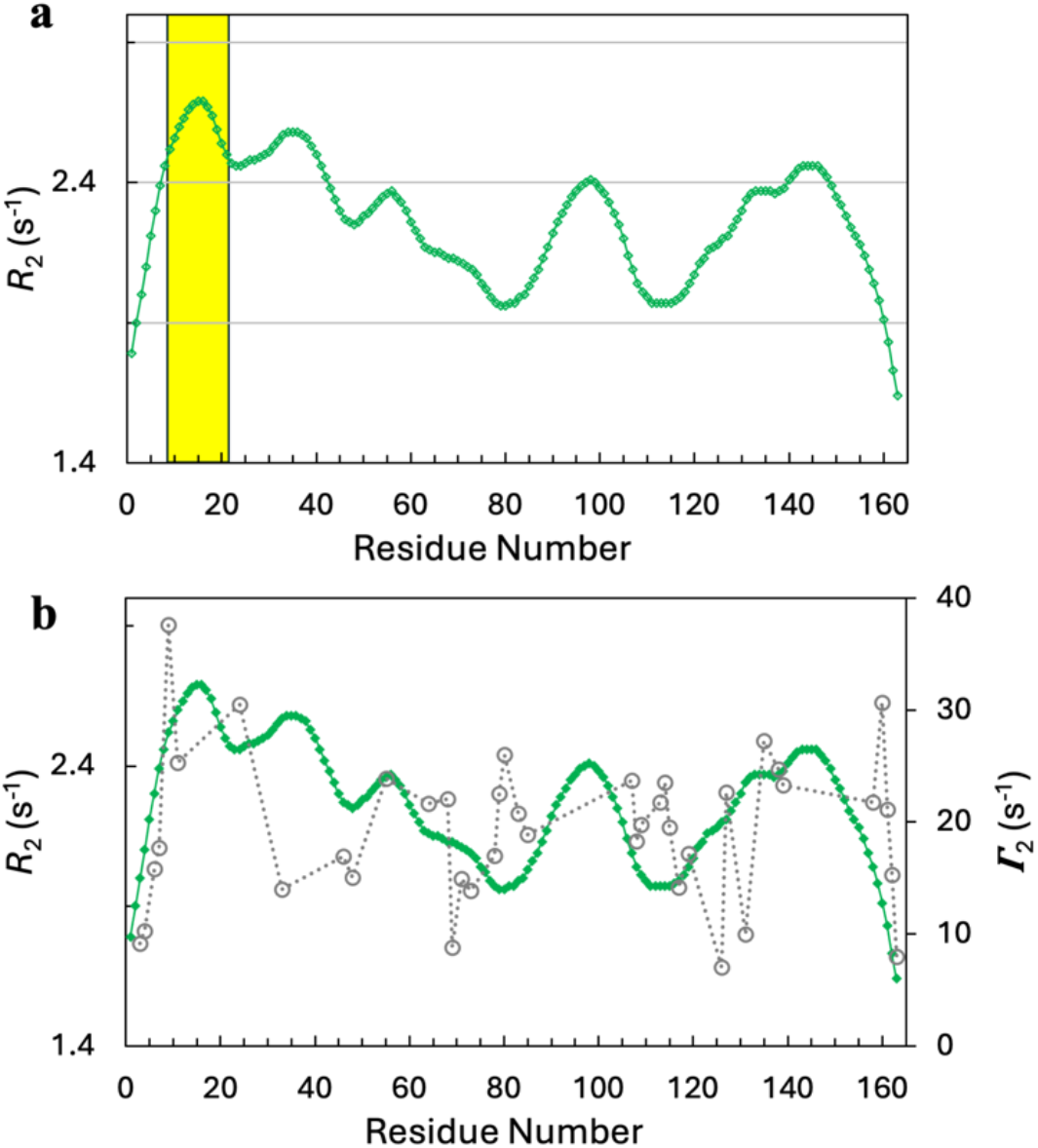
SeqDYN predicts the sequence motif driving the phase separation of FUS. (a) *R*_2_ profile. Yellow shading highlights the highest peak, spanned by Q9-P21 that was supported by PRE effects ^28^. (b) Overlay of the *R*_2_ profile and the *Γ*_2_ profile generated by a spin label attached to residue 16.

Wang et al. ^10^ observed a strong correlation between the threshold concentration for phase separation and the numbers of Tyr and Arg residues among the FUS family of proteins. These authors further studied the contributions of Tyr and Arg residues in FUS phase separation by making Y-to-F, R-to-K, polar-to-R, and other substitutions, mostly in the disordered N-terminal 40% of the sequence (residues 1-211) but also in the remainder of the sequence (residues 212-526). As already noted, Tyr and Arg are in the top tier of amino acids associated with elevated *R*_2_ rates. We actually found a strong correlation between − In(*C*_th_) and the mean *R*_2_ of the whole sequence (*r* = 0.85; Figure S2a). A single outlier is the Q-to-G mutant; the threshold concentration is minimally perturbed but the SeqDYN-predicted *R*_2_ profile experiences a large decrease in mean value. MD simulations have shown that Gly play dual roles in IDP dynamics: it has minimal interactions with other amino acids (hence often cited as a “spacer”) and thus lowers *R*_2_, but its flexibility also facilitates interactions between other amino acids and thereby can lead to elevated *R*_2_ ^36^. The latter role is not modeled by SeqDYN.

### DDX4: clustering of interaction-prone amino acids along the sequence

Brady et al. ^27^ reported intermolecular NOEs in the dense phase of the DDX4 N-terminal IDR (residues 1-236), showing the importance of Phe and Arg in driving phase separation. The SeqDYN-predicted *R*_2_ profile exhibits five major peaks, at I_20_F_21_, H_53_F_54_, F_109_WR_111_, F_127_S_128_, R_146_R_147_ (Figure 5). Each of these short motifs contains either a Phe or an Arg residue or both. These short motifs may be centers of local clusters that mediate phase separation. Indeed, sequence analysis of DDX4 orthologs by Nott et al. ^8^ revealed that Phe and Arg residues are clustered along the sequence. Moreover, like charges are also clustered along the sequence, and charge scrambling resulted in ablation of LLPS.

**Figure 5.**
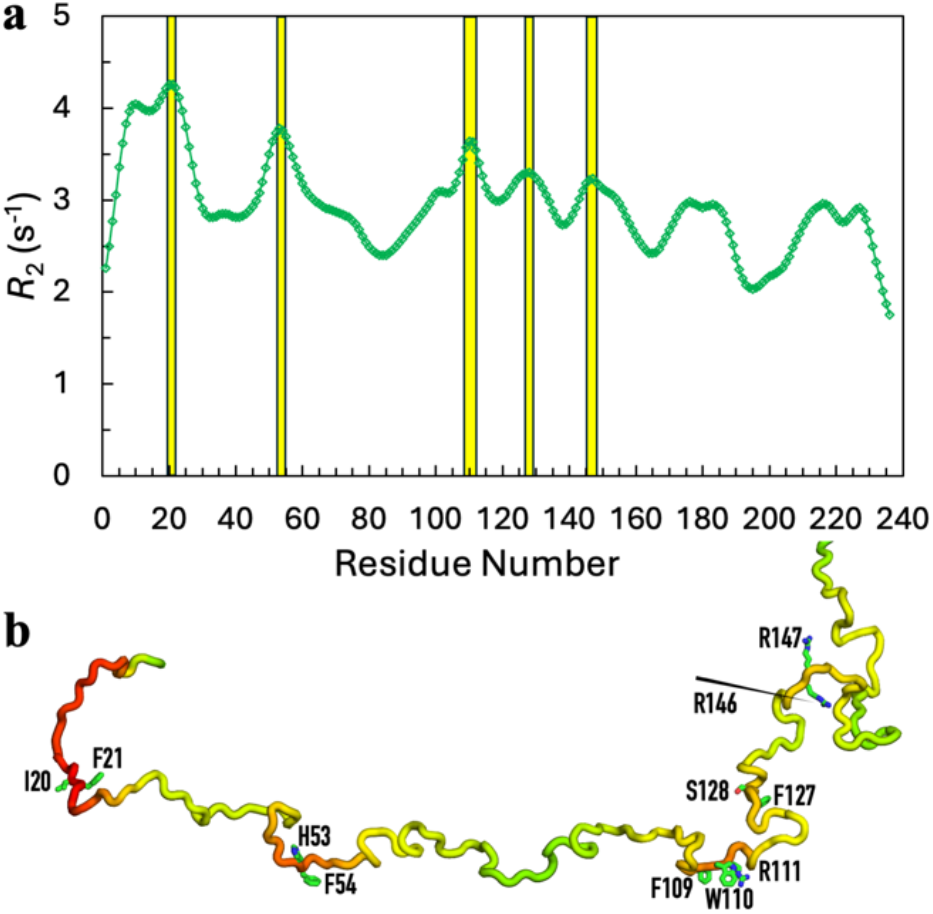
SeqDYN predicts the sequence motifs potentially driving the phase separation of DDX4. (a) *R*_2_ profile. Yellow shading highlights five major peaks at I_20_F_21_, H_53_F_54_, F_109_WR_111_, F_127_S_128_, R_146_R_147_, which contain Phe and Arg that were shown to be important for phase separation by NOESY ^27^. (b) *R*_2_ displayed as a color spectrum (green-yellow-orange-red) rendered on a representative DDX4 structure. Sidechains are displayed for motifs highlighted in (a).

Recently Norrild et al. ^41^ used partition of DDX4 fragments in DDX4 condensates to probe sequence motifs that drive phase separation. By averaging over overlapping fragment tiles, they obtained a partition profile along the sequence, which putatively reflects the sequence-dependent drive for phase separation. The partition profile exhibits a periodicity of ∼40 residues. This periodic pattern is captured well by the SeqDYN-predicted *R*_2_ profile (Figure S3a). The valleys can be attributed to the clustering of low-*q* amino acids including Gly and Ser along the sequence, such as in the segment S_80_TMGGFGVG_88_, whereas the peaks can be attributed to the clustering of high-*q*, i.e., interaction-prone, amino acids including Phe and Arg. Indeed, Norrild et al. found that the level of partition in DDX4 condensates increased with the number of aromatic residues in the fragment.

These authors found that a composition-based model can explain their partition data to a large extent, but including a position-dependent contribution narrowed the gap between model and data. They further showed, after scrambling the sequences of 16-residue tiles, the partition data showed good agreement with the counterparts from the tiles with WT sequences. The good performance of the composition-based model and the good match in partition data between WT and scrambled tiles led to the conclusion that amino-acid composition, rather than sequence motifs, is the main drive of DDX4 phase separation. However, *R*_2_ prediction by SeqDYN suggests a different perspective. There is no question that amino-acid composition is a major determinant for phase separation, as all the afore-mentioned studies have noted the importance of aromatic residues. The four aromatic residues (Trp, Tyr, Phe, and His) are among the top-6 according to both the *q* parameters in SeqDYN and the free energy contributions in Norrild et al.’s composition-based model. Indeed, there is a strong correlation (*r* = 0.79; Figure S3b) between these two sets of parameters among the 20 types of amino acids. The issue in contention is whether, for the given composition of an IDP, the positions of the amino acids along the sequence also play a crucial role. Norrild et al.’s own data and those of Nott et al. ^8^ support a positive answer. As stated above, adding a position-dependent contribution more fully explained the partition data. Also, residues 132-166 that span a peak both in the partition profile and the *R*_2_ profile (Figure S3a) ablated phase separation ^8^, suggesting that some sub-regions may be particularly important.

A correlation between the partition profiles of WT and scrambled tiles may appear to support the unimportance of amino-acid positions. However, recall that the correlation half-length *L*_1/2_ has a value of 5.6, meaning that a single correlated segment can extend to 11 residues or longer. In another words, a correlated segment, rich in high-*q* residues, will still be a correlated segment even after scrambling. Likewise, a segment that is rich in low-*q* residues will still be rich in low-*q* residues after scrambling and thus appears uncorrelated. I illustrate this point by comparing the *R*_2_ profile of the WT DDX4 sequence with those of two variants, each with a 16-segment scrambled (Figure S3c). One segment, residues 11-26, spans a peak in the WT *R*_2_ profile; after scrambling, an *R*_2_ peak still appears over this segment, with an elevation in peak height and a slight shift in peak position. The second segment, residues 81-96, spans a valley in the WT *R*_2_ profile; after scrambling, an *R*_2_ valley still appears over this segment, with a smal shift in valley position. This illustration explains the correlation between the partition profiles of WT and scrambled tiles.

While the *R*_2_ profile overall tracks well the partition profile, there is a notable underestimation of the peak height around R150 (Figure S3a). DDX4 has a net charge of −4, whereas the segment around R150 is enriched in basic residues (net charge +4 for residues 146-154). The high partition peak around R150 can thus be attributed to electrostatic attraction. SeqDYN only models local interactions and thus misses long-range (or inter-chain) electrostatic attraction. In the present case, the latter effect can be mimicked by enlarging the *q* values of basic residues and suppressing the *q* values of basic residues, which significantly improves the agreement between the partition and *R*_2_ profiles (Figure S3a).

### A1-LCD: rule or exception?

A1-LCD (residues 186-320 of Uniprot P09651-2) presents a prime example for the idea that IDPs use the composition, rather than the positions, of amino acids for driving phase separation ^12^. This IDP has a high aromatic content (14% of the sequence). These aromatic residues are more uniformly distributed than 99.99% of randomly generated sequences. Correspondingly, the SeqDYN-predicted *R*_2_ profile is relatively unform, except for a peak around residue K115, which is close to two consecutive aromatic residues, Y112 and F113 (Figure 6a). When the aromatic content was increased by 1/3, the critical temperature (*T*_c_) for phase separation increased; conversely, when the aromatic content was decreased by 1/3 and 2/3, *T*_c_ decreased successively ^12^. The increase in aromatic content leads to elevation in *R*_2_ around the peak at K115 and new peak around R11, whereas the decreases in aromatic content lead to largely uniform reduction in *R*_2_. The mean *R*_2_ values of these variants show a perfect correlation with the *T*_c_ values (Figure 6b).

**Figure 6.**
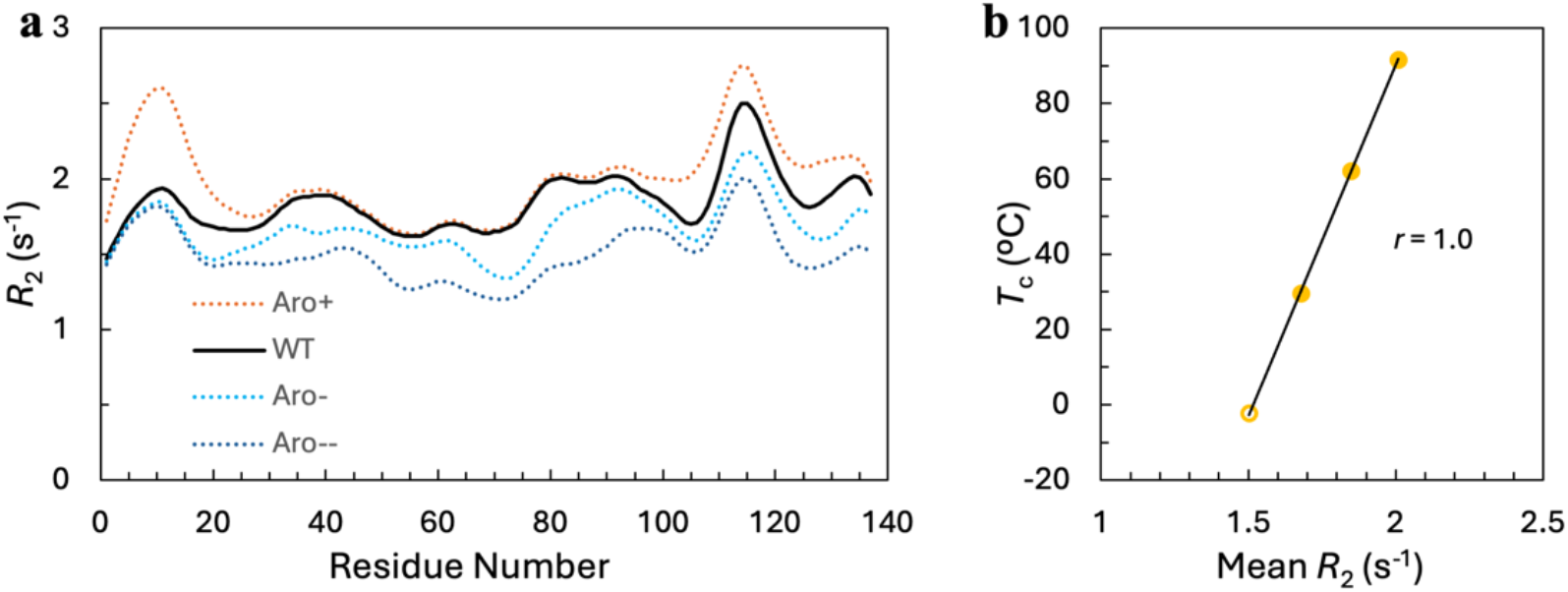
SeqDYN quantitatively accounts for the effects of aromatic mutations on the critical temperature of A1-LCD phase separation. (a) *R*_2_ profiles of WT A1-LCD and three aromatic mutants. (b) Correlation between *T*_c_ and mean *R*_2_. *T*_c_ from binodals in Martin et al. ^12^.

When the aromatic residues of A1-LCD were repositioned into clusters of three or four along the sequence, the mutant formed amorphous aggregates instead of liquid droplets, suggesting overly strong attractions ^12^. So one interpretation of the highly uniform aromatic distribution in the A1-LCD sequence is that the high aromatic content by itself also produces sufficient attraction for LLPS; clustering of these aromatic residues will increase the drive for phase separation, but at the risk of producing overly strong attraction as to resulting in solid-like aggregates instead of liquid droplets.

Martin et al. ^12^ and Bremer et al.^14^ measured the threshold concentrations of over 30 A1-LCD mutants. The latter authors observed an increase in *C*_th_ with increasing net charge, but the minimum in *C*_th_ occurs a net charge of not 0 but 3 to 4. There is a good correlation between − In(*C*_th_) and a variable that combines the mean 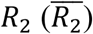 and the net charge (*Q*):

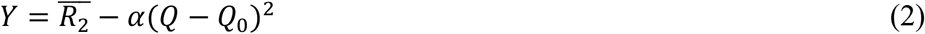

(*r* = 0.60; Figure S2b), with *Q*_0_ = 4 and *α* = 0.01. Again, a mutant involving 23 S-to-G substitutions is an outlier. Overall, Figures S2 and 6b demonstrate that SeqDYN can quantitatively model the effects of mutations on the phase-separation propensities of IDPs.

In contrast to the nearly uniform distribution of aromatic residues along the sequence of A1-LCD, Johnson et al. ^15^ found that the clustering of Tyr residues promotes the phase separation of the LCD (residues 1-264) of the RNA-binding protein EWS according to NMR data and mutational effects. The NMR data included backbone ^15^N transverse relaxation rates, which enable a direct test of SeqDYN predictions. As shown in Figure S4a, SeqDYN predictions match well with the experimental data, except for a mild underestimation in the C-terminal region. The mean *R*_2_ values of SeqDYN also model well the phase-separation propensities of WT EWS-LCD and six Y-to-S and one N-to-S mutant as measured by pelleting assays (*r* = 0.91; Figure S4b).

### Correlated segments driving LLPS of other IDPs

Using deletion constructs, Newton et al. ^34^ found that residues 1-152 are responsible for the phase separation of LINE-1 ORF1 protein. Residues 1-52 are disordered but residues 53-152 can form a coiled-coil. Based on further deletions and *R*_2_ measurements of the ORF1 1-152 construct and of the 1-52 fragment with and without adding the 53-152 fragment, the authors concluded that the interactions of residues 47-53 and 132-152 are required for the phase separation of the ORF1 1-152 construct. The SeqDYN-predicted *R*_2_ profile is presented in Figure 7a, where residues 47-53 are at the middle of a stretch with elevated *R*_2_ while residues 132-152 span the highest peak. Newton et al. reported *R*_2_ data for the first 50 residues of the ORF1 1-152 construct, which show a rapid rise around residue 30. This trend is accurately predicted by SeqDYN. The SeqDYN-predicted *R*_2_ profile has one additional peak, around residue R98; a mutation at residue 93 enhanced the phase-separation propensity, suggesting that residues spanning this peak may also contribute to the drive for phase separation.

**Figure 7.**
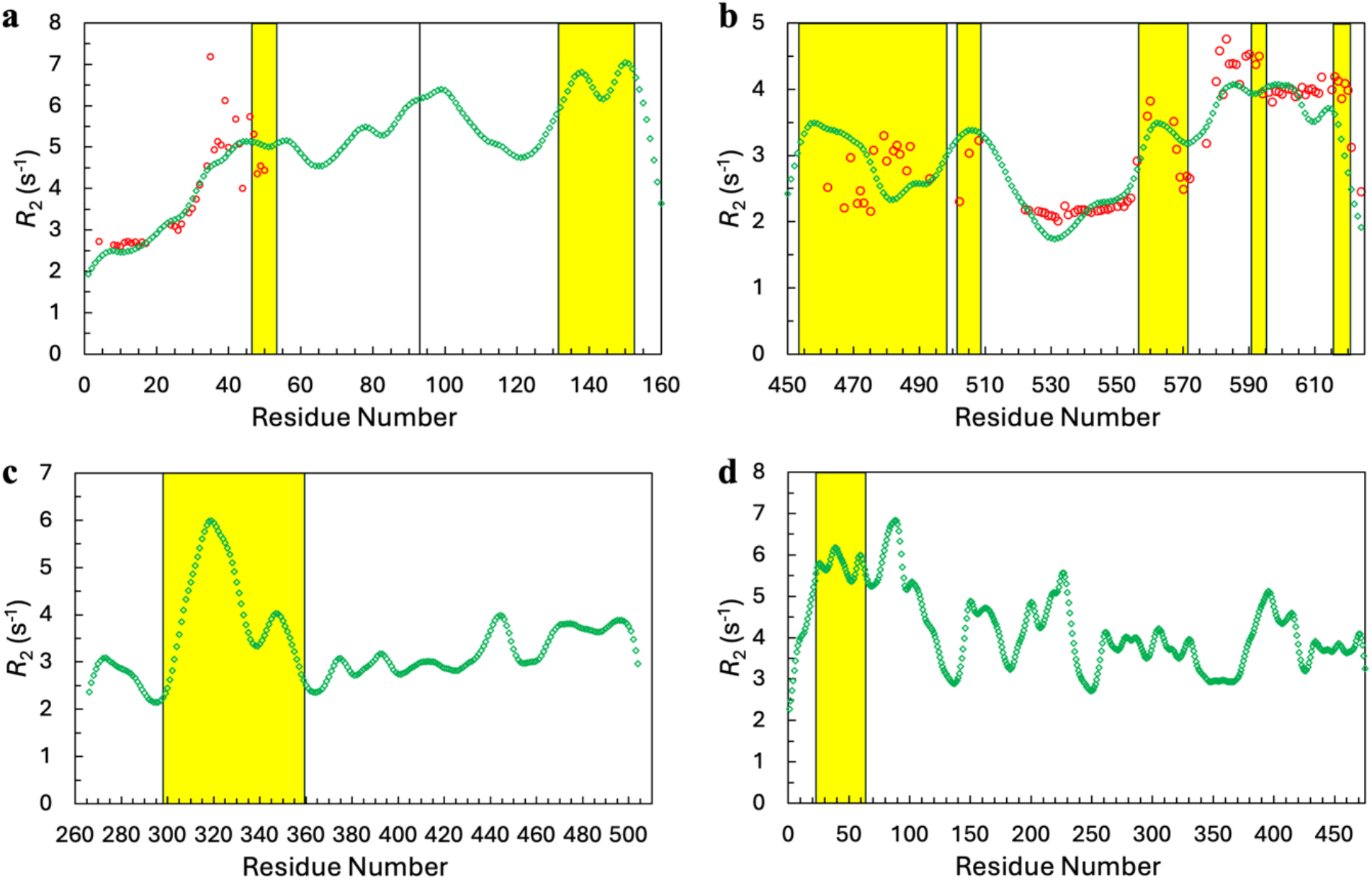
SeqDYN predicts the sequence motifs and sub-regions driving the phase separation of four other IDPs. (a) The ORF1 1-152 construct. Green diamonds display SeqDYN predictions whereas red circle display experimental data of Newton et al. ^34^. These authors concluded that the interactions of residues 47-53 and 132-152, highlighted by yellow shading, were required for LLPS. A mutation of residue 93 (vertical line) enhanced LLPS. (b) The UBQLN2 450-624 construct. Green diamonds display SeqDYN predictions whereas red circle display experimental data of Dao et al. ^33^. These authors concluded that interactions of five sub-regions, highlighted by yellow shading, drive phase separation. (c) The YAP 266-504 IDR. A coiled-coil sub-region, highlighted by yellow shading, is required for LLPS s ^26^. (d) U2AF^65^. The RS region, highlighted in yellow, drives LLPS ^25^.

Using a similar deletion approach, Dao et al. ^33^ identified residues 450-624 as the minimal construct for the phase separation of proteasomal shuttle factor UBQLN2. Based on concentration-dependent chemical shift perturbations and elevated *R*_2_ rates, Dao et al. ^33,42^ concluded that interactions of residues 454-498, 502-508, 557-572, 591-595, and 617-622 drive oligomerization and phase separation of the UBQLN2 450-624 construct. SeqDYN predictions track the experimental *R*_2_ well and feature these interaction motifs as major peaks (Figure 7b).

The C-terminal IDR (residues 266-504) of YAP phase separated in vitro and in cells ^26^. Based on deletion constructs, a coiled-coiled sub-region (residues 298-359) was found to drive the LLPS. This sub-region stands out in the SeqDYN-predicted *R*_2_ profile as it spans the single, tall peak (Figure 7c). Likewise, the RS region (residues 23-63) drives the phase separation of U2AF^65^ ^25^. This region corresponds to a major peak in the SeqDYN-predicted *R*_2_ profile (Figure 7d). An even taller peak occurs around residue R89. Whether residues there also contributes to LLPS remains to be tested.

## Discussion

SeqDYN is able to rationalize a variety of experimental observations on the phase separation of IDPs. Previous studies have identified sequence motifs that drive phase separation, including G624–R626, G638–R640, and R660–Q666 in CAPRIN1 by NOESY ^32^, R21-G30 in LAF-1 by coarse-grained simulations and deletion ^13^, and Q9-P21 in FUS by PRE ^28^. All these motifs manifest themselves as major peaks in SeqDYN-predicted *R*_2_ profiles, which can be attributed to the clustering of amino acids that strongly interact with each other. Moreover, the *q* parameters of SeqDYN strongly correlate with the threshold concentrations of XXssXX peptides, and the changes in the mean *R*_2_ value of the entire sequence quantitatively model the effects of mutations on the threshold concentrations and critical temperatures of IDPs. Lastly, SeqDYN-predicted *R*_2_ profiles closely track partition profiles of fragment tiles inside IDP condensates.

Empirical formulas have been proposed to model phase-separation properties using only amino-acid composition ^10,14,17-19,41^. SeqDYN departs from these models by accounting for the effects of interaction clusters. It is also notable that SeqDYN was parameterized using single-chain data, i.e., the backbone ^15^N transverse relaxation rates of 45 IDPs, and originally was not even designed for predicting phase-separation properties. As more and more sequence-specific phase-separation data become available, it will be possible to train a SeqDYN-style model on such data. I have already illustrated that SeqDYN can be tailored to account for charge effects [Figure 3a and eq (2)]. The dual effects of Gly and potential effects of residual secondary structures will deserve special attention.

A1-LCD typifies the view of amino-acid composition as the driver of homotypic IDP phase separation. In this view (Figure 8a), stickers are disbursed along the sequence and sticker-sticker interactions rarely form within a single chain in the dilute phase. In the dense phase, IDP chains cross each other; at each crossing, the inter-chain interaction is mostly formed by a pair of sticker residues. For many IDPs, these pairwise interactions may be too weak for forming a stable dense phase. Instead, a motif-based view may be posited. In this view (Figure 8b), stickers are clustered, leading to correlated segments, along the IDP sequence. In the dilute phase, these segments rarely interact with each other within a single chain. In the dense phase, correlated segments from different chains interact with each other, with intra-chain pairs broken up to form inter-chain pairs. Correlated segments play similar roles as charge blocks and as interaction motifs that target folded domains (e.g., SH3 and SUMO), and may be the drivers of phase separation for many IDPs.

**Figure 8.**
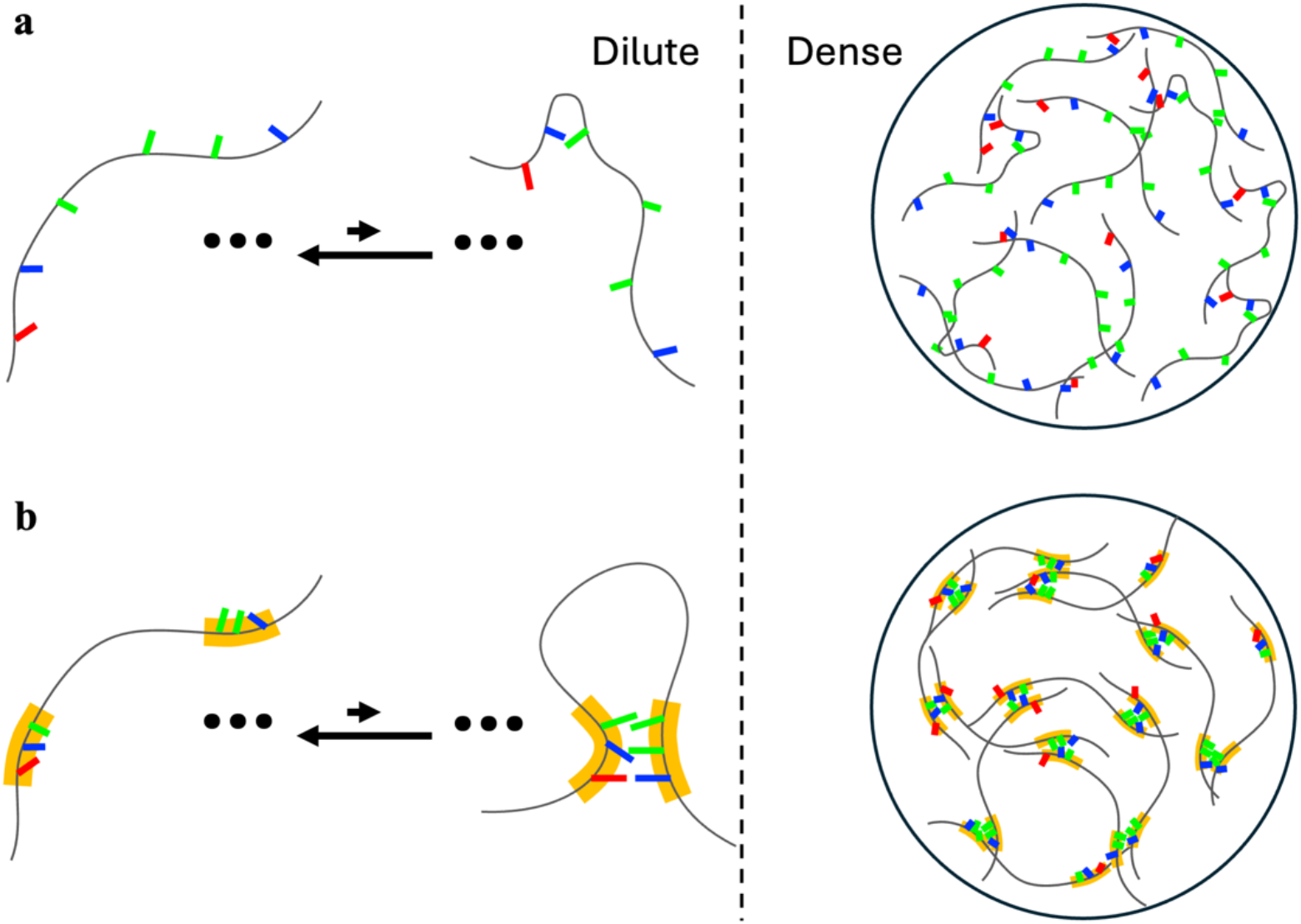
Two views on the drivers of IDP phase separation. (a) Amino-acid composition based view. (b) Correlated-segment based view.

In conclusion, ample evidence suggests that the phase separation of many IDPs may be driven by clusters of interaction-prone amino acids along the sequence, rather than the amino acids separately. SeqDYN predicts these clusters as correlated segments that span major peaks in the *R*_2_ profile. There is a limit to what can be accounted for by amino-acid composition alone; the positions of amino acids along the sequence provide an additional handle for dissecting the drive for IDP phase separation and for designing sequences with desired phase-separation properties.

## Computational Methods

All SeqDYN predictions were obtained using the web server at https://zhougroup-uic.github.io/SeqDYNidp/ ^37^. To account for DDX4-fragment electrostatic interaction (Figure S3a), the *q* values of Arg and Lys were enlarged by 1.1-fold whereas those of Glu and Asp were reduced by 1.1-fold. To account for the effect of net charge on the threshold concentration of A1-LCD (Figure S2b), we combined the mean *R*_2_ with the net charge via eq (2).

## Acknowledgment

This work was supported by National Institutes of Health Grant GM118091.

