## Supplementary Figures for "Correlated Segments of Intrinsically Disordered Proteins as Drivers of Homotypic Phase Separation"

Huan-Xiang Zhou*

Department of Chemistry and Department of Physics, University of Illinois Chicago, Chicago, IL 60607, USA

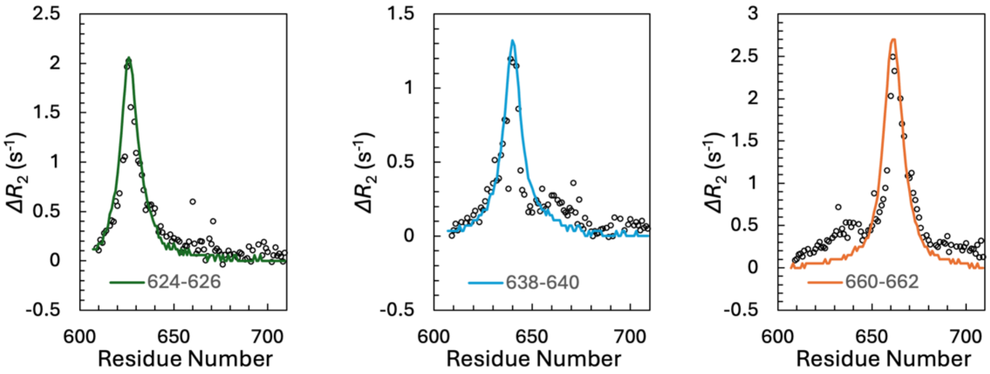


Figure S1. SeqDYN accurately predicts the changes in $R_2$ profile upon mutating three consecutive residues into ASA. $\Delta R_2$ is WT minus mutant; mutated residues are indicated in legends. Symbols are NMR data from Kim et al. [32]; curves are SeqDYN predictions.

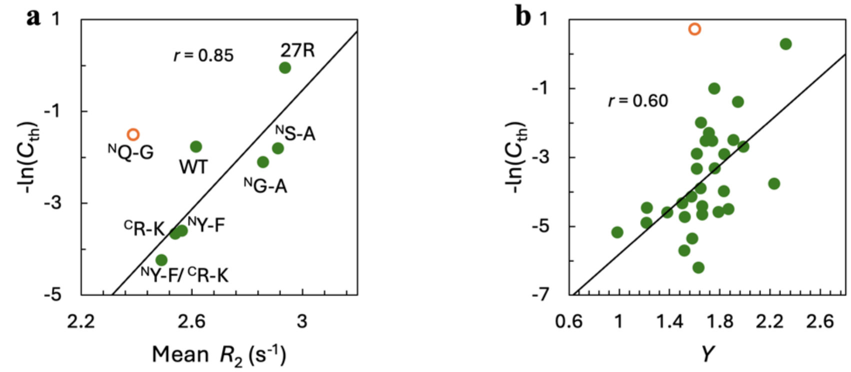


Figure S2. SeqDYN quantitatively accounts for the effects of mutations on the threshold concentrations. (a) Correlation between $-\ln(C_{\mathrm{th}})$ and mean $R_2$ for full-length FUS. $C_{\mathrm{th}}$ data at 150 mM KCl from Wang et al. [10]. "N" and "C" indicate N-terminal or C-terminal regions. (b) Correlation between $-\ln(C_{\mathrm{th}})$ and the compound variable $Y$ for A1-LCD. $C_{\mathrm{th}}$ data from Martin et al. [12] and Bremer et al. [14], as tabulated by Ibrahim et al. [17].

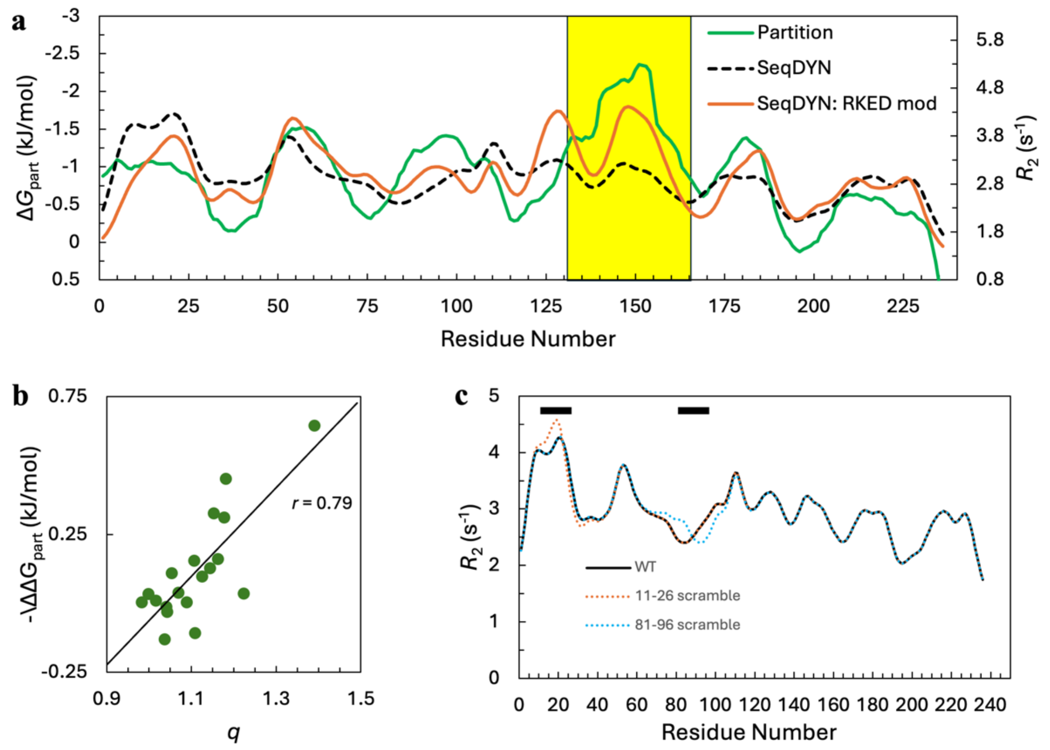


Figure S3. SeqDYN-predicted $R_2$ profile matches well with the partition profile of fragment files in DDX4 condensates. (a) Overlay of the $R_2$ profile with the partition profile. Partition data from Norrild et al. [41]; "RKED mod" means SeqDYN prediction with $q$ values enlarged for Arg and Lys and suppressed for Glu and Asp. (b) Correlation between amino-acid contributions to partition free energy and $q$ parameters. (c) Effects of sequence scrambling within 16-residue segments on predicted $R_2$ profile. These two segments are indicated by black bars at the top.

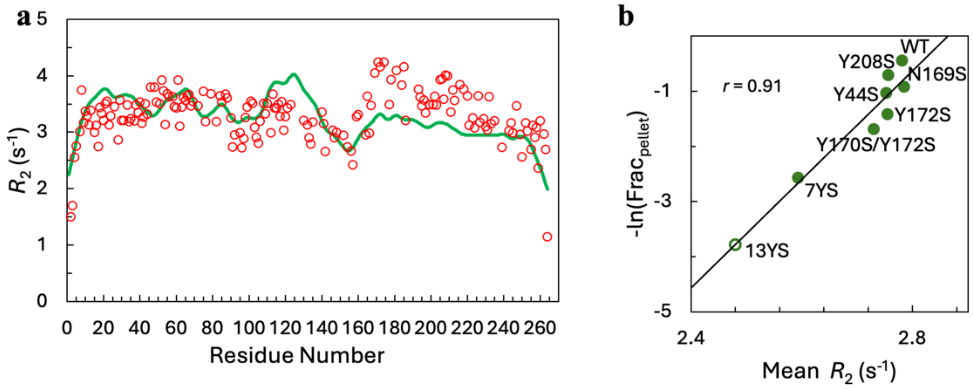


Figure S4. SeqDYN accurately predict the $R_2$ profile of the EWS LCD and the phase-separation propensities of eight variants. (a) Predicted (green diamonds) and experimental (red circles [15]) $R_2$ profiles. (b) Correlation between reported pellet fractions [15] and SeqDYN mean$R_2$ values. Correlation analysis was performed on seven variants; a very low pellet fraction (open circle) predicted for the eight variant, 13YS, is consistent with the reported undetectable pellet fraction.
